# Scaling Logical Density of DNA storage with Enzymatically-Ligated Composite Motifs

**DOI:** 10.1101/2023.02.02.526799

**Authors:** Yiqing Yan, Nimesh Pinnamaneni, Sachin Chalapati, Conor Crosbie, Raja Appuswamy

**Author notes:** Contributing authors.

## Abstract

DNA is a promising candidate for long-term data storage due to its high density and endurance. The key challenge in DNA storage today is the cost of synthesis. In this work, we propose *composite motifs*, a framework that uses a mixture of prefabricated motifs as building blocks to reduce synthesis cost by scaling logical density. To write data, we introduce Bridge Oligonucleotide Assembly, an enzymatic ligation technique for synthesizing oligos based on composite motifs. To sequence data, we introduce Direct Oligonucleotide Sequencing, a nanopore-based technique to sequence oligos without assembly and amplification. To decode data, we introduce Motif-Search, a novel consensus caller that provides accurate reconstruction despite synthesis and sequencing errors. Using the proposed methods, we present an end-to-end experiment where we store the text “HelloWorld” at a logical density of 84 bits/cycle (14–42**×** improvement over state-of-the-art.)

## 1 Introduction

The growing adoption of Big Data Analytics and Artificial Intelligence has led to an explosion in the rate of data generation. A recent survey by the International Data Corporation reports that the digital datasphere is forecast to grow to 125 zettabytes by 2025 [1] and is anticipated to exceed silicon supply in 2040 [2]. As traditional storage media is unable to keep pace with the rate of data growth [3], synthetic DNA has become an attractive archival storage medium due to its high density, longevity, and absence of technical obsolescence compared with electronic media [4–8].

In most prior work on DNA-based digital storage [5, 7, 9, 10], DNA synthesis is based on phosphoramidite chemistry [11], a technology that has been optimized over several decades to perform highly-accurate, base-by-base synthesis of short DNA strands by making phosphodiester bonds between nucleotides. There are three Key Performance Indicators (KPIs) that can be used to evaluate the efficiency of DNA synthesis: (i) bits written per cycle (also called *logical density* [12, 13]), (ii) bits written per oligo, and (iii) coupling reactions per oligo. The efficiency of writing data to DNA depends on the number of synthesis cycles (*x*) to grow the strand and available repeating units (*m*) for addition at each cycle. The information capacity of the oligo (*N* bits) can be derived as

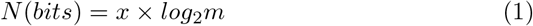

While base-by-base synthesis methods can perform 200 or more coupling cycles(*x*), the number of available subunits to add at each cycle is four (nucleotides), thereby limiting bits per synthesis cycle to two, and the information capacity of an oligo to a few hundred bits. While the quality, quantity, cost, and rate of DNA synthesis provided by base-by-base chemistry is suitable for biological research, it is far from ideal for the DNA storage use case. This has resulted in synthesis emerging as a major bottleneck in DNA storage.

In this work, we introduce the *composite motifs* framework to scale logical density well beyond the limit of 2 bits per synthesis cycle. Composite motifs are inspired by recent advances in motif-based approaches to DNA data storage [14, 15] that use short oligonucleotide sequences, also referred to as motifs, that are drawn from a fixed library as building blocks for assembling longer oligos. Using a motif library of *M* motifs, one can scale logical density by storing *log*_2_(*M*) data bits per synthesis cycle. The use of a fixed library of motifs similar to a typesetting press can also simplify miniaturization and automation. The composite motif framework builds on the benefits of motif-based DNA storage, and further improves logical density by exploiting sequencing multiplicity inherent in DNA synthesis by encoding data using a combination of motifs rather than individual motifs.

In this work, we show that a DNA storage system based on composite motifs can provide an order of magnitude improvement in logical density over state-of-the-art systems by implementing an end-to-end prototype system. In doing so, we develop new encoding and enzymatic motif ligation techniques that can scale DNA synthesis in the DNA write pipeline, and assembly-free, Nanopore-based motif read out and alignment-based motif decoding techniques that can scale DNA sequencing in the DNA read pipeline.

## 2 Results

### 2.1 Composite Motifs as Building Blocks for DNA Storage

A composite motif is a representation of a position in an oligo sequence that uses a combination of motifs drawn from a fixed motif library to encode data. For example, assuming a library of 32 motifs, and a combination factor of four, there are *C*(32, 4) = 35960 possible unique combinations with which we can encode 15 (*log*_2_35960) bits of data per composite motif. Composite motifs increase logical density by expanding the motif library using combinations of motifs without increasing the volume of motifs. As current synthesis platforms already use a high degree of sequence multiplicity (multiple copies of DNA molecules are synthesized per oligo), composite motifs can also be integrated into current platforms without any extra cost as they can exploit sequence multiplicity to scale logical density. Higher logical density also leads to a reduction in the length of DNA required to store the same amount of data, alleviating issues related to long oligo synthesis.

In order to demonstrate the feasibility of using composite motifs, we developed a DNA storage system that uses composite motifs as building blocks. Fig 1 presents the read/write pipeline of our system. On the writing side, digital data is encoded into oligos containing composite motifs using a motif encoder. Writing a composite motif at any given position of an oligo sequence is done by mixing multiple motifs during the synthesis procedure to synthesize multiple DNA molecules that contain the corresponding combinations of motifs using *Bridged Oligonucleotide Assembly* (BOA) (Sec.2.2). On the reading side (Fig 1), we read composite motifs by amplification-free sequencing of multiple DNA molecules using *Direct Oligonucleotide Sequencing* (DOS) (Sec.2.3), and then decode the data using our new motif-based consensus caller called *Motif-Search*(Sec.2.5).

**Fig. 1.**
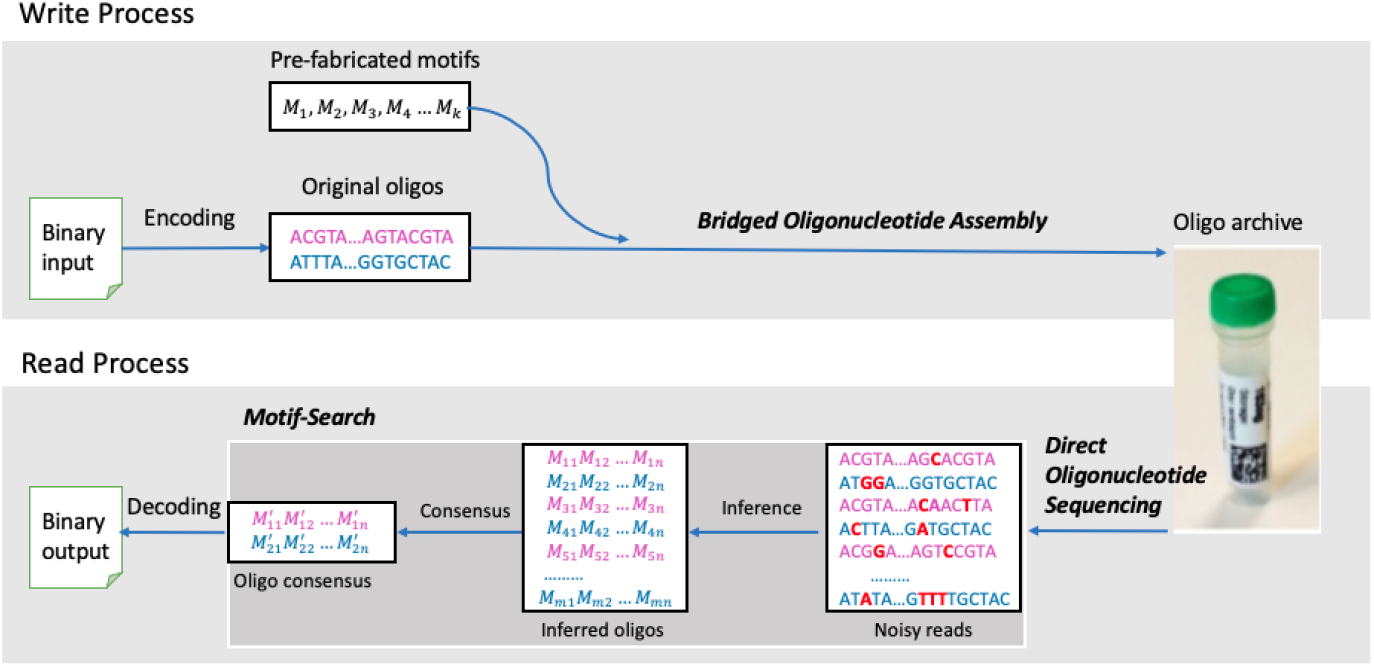
Composite-motif-based data write and read pipeline.

### 2.2 Bridged Assembly of Composite Motifs

#### Encoding

In order to demonstrate the feasibility of composite motifs, we stored the text “HelloWorld” using our composite-motif-based DNA storage system. The sequence design rules for base motifs that are used to derive composite motifs are similar to those of DNA barcode design. Thus, we started with DNA sequences designed in prior work [16] to select 96 25nt base motifs. Using a combination factor of 32, we developed a composite motif set of 3 × 10^25^ composite motifs (*C*(96, 32)). Thus, each composite motif, and hence, each synthesis cycle, can store 84-bits of data (*log*_2_*C*(96, 32)). As our input text is 10 bytes, it can be stored using a single DNA sequence with one composite motif. However, in order to test precision and recall of methods in the read pipeline, we stored the same data eight times using eight sequences. We index each sequence using eight unique 24nt address motifs that are separate from the 96 payload motifs. While we designed our encoding to match our experimental needs, it naturally extends to a general purpose encoder(Fig 2.a).

**Fig. 2.**
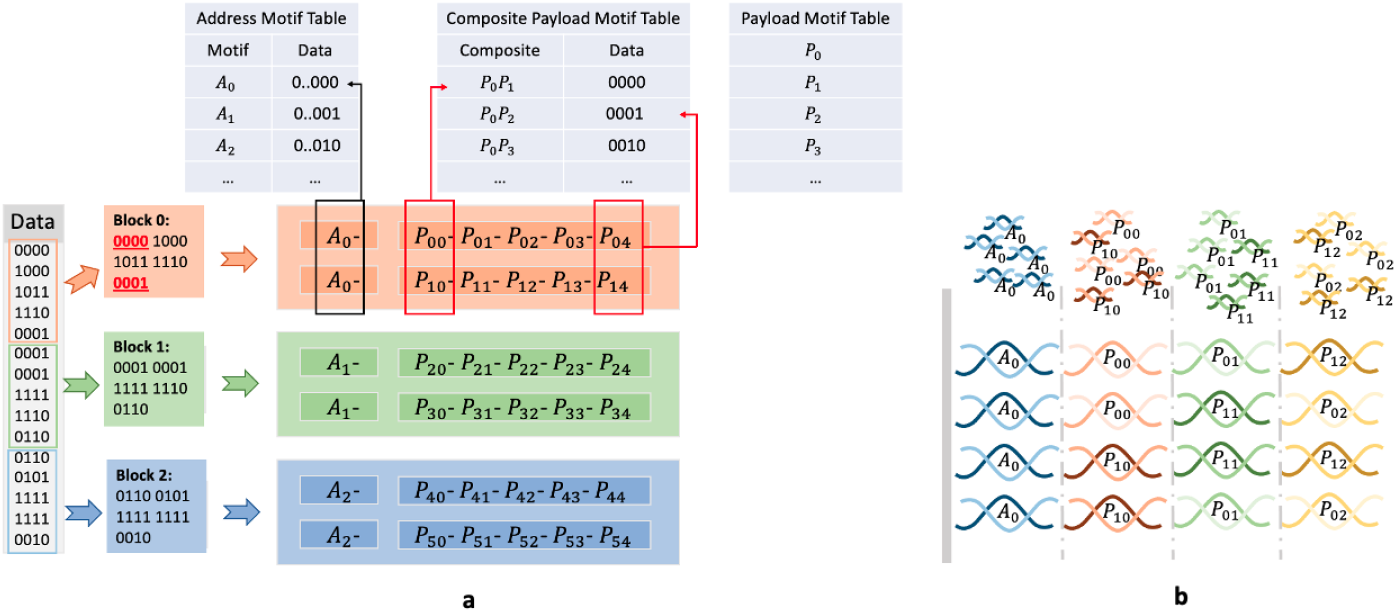
Composite motifs increases the logical density in DNA-based storage. (**a**) A block of binary data is encoded to a sequence comprising a set of oligos with same address payload motifs. The composite of payload motifs from the same vertical position represents the binary data together. (**b**) Composite motifs can be generated by mixing the motifs during each synthesis cycle. A: address motif, P: payload motif.

#### Bridge Oligonucleotide Assembly

Each of the eight encoded sequences is then used to synthesize 32 oligos producing a total of 256 oligos. The address motif is repeated in each molecule, while the composite motif is expanded to generate a variant combination using 32 payload motifs. Oligos are synthesized using template-directed ligation. This method utilises single-strand sequences, referred to as *bridge* oligos, to facilitate the ligation of payload motifs to address motifs. In the general case, an oligo would contain one or more address and payload motifs as shown in Fig 2. As any motif can be ligated with any other, designing bridge oligos for each possibility is suboptimal and not scalable. We solve this problem by using a *spacer* motif. When the motif library is designed, each 25nt motif is extended on both 5’ and 3’ ends with 12nt and 13nt nucleotides from the 3’ and 5’ ends of the spacer motif (Fig 3.a). While this increases the length of each synthesized motif from 25nt to 50nt, it does not affect the number of motifs, and more importantly, it makes it possible to design the bridge oligo to be complementary to a single spacer. By doing so, the bridge oligos can hybridise to the spacer portions at the 3’ and 5’ ends of two payload motifs while the enzyme ligates them.

**Fig. 3.**
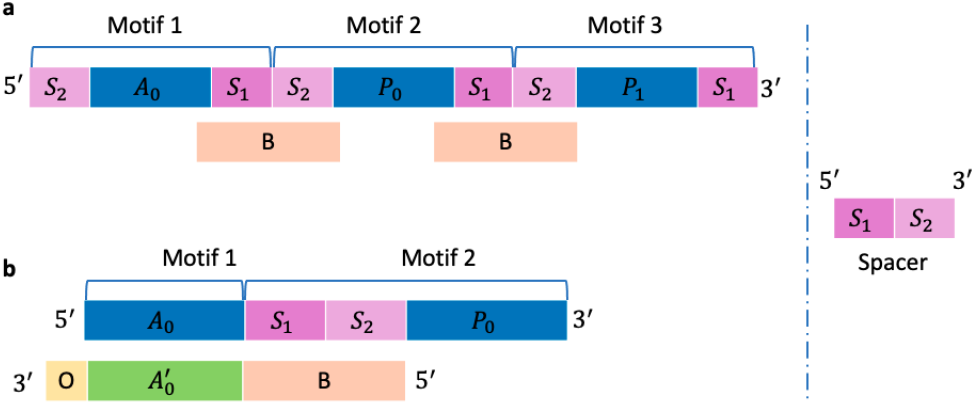
Bridged oligonucleotide assembly. (**a**) The general oligo structure design. (**b**) The experimental oligo structure design. A: address motif, A’: reverse complement of A, P: payload motif, S: spacer, B: bridge, O: overhang.

For the purpose of our experiment, as we have only 2 motifs per oligo, we modified this by (i) prepending the entire spacer sequence to the 5’ end of each payload motif, and (ii) designing eight (instead of one) bridge oligos, each of which is complementary to both the spacer sequence and one of the eight address sequences (Fig 3.b). By doing so, the eight bridge motifs also double in role as adapters during sequencing. The spacer-extended 32 payload motifs, eight address motifs, and eight bridge oligos were all synthesized base-by-base by Integrated DNA Technologies (IDT). The oligos were synthesized by selecting, annealing and ligating together the corresponding address–payload motif pairs. The inputs to the reaction comprise all motif oligos, bridge oligos, enzymes and ligation buffer. These reactions proceeded to produce ligated oligos through programmed temperature incubation and cycling, where each bridge oligo facilitates the ligation of a specific address motif with a payload motif via complementary annealing. We use the resulting oligo pool to test the feasibility of decoding the identity of motifs from an enzymatically-ligated, Nanopore-basecalled readout.

### 2.3 Direct Nanopore Sequencing & Error Characterization

A key aspect of a DNA storage system, along with DNA writing performance, is the cost of DNA sequencing and time taken to read data from DNA molecules. Nanopore sequencing enables single molecule sensing capabilities and has the potential to create a low-cost, high-speed DNA storage read head. The yield of a Nanopore (ONT) flowcell is dependent on the size of the DNA to be sequenced. Small oligos result in a higher number of unoccupied pores over time. ONT estimates that the minimum DNA size to load in a R9.4 flowcell is 200 bases. Thus, prior work on DNA storage with Nanopore has relied on additional sample preparation steps for short oligos that are manual and time consuming [17]. In particular, DNA assembly methods were used to concatenate five or more DNA storage oligos into a longer fragment, and PCR amplification is used to sufficiently increase sequencing throughput and coverage for decoding.

We developed a method to enable direct sequencing of composite-motif-based oligos without amplification or second-strand synthesis. As mentioned earlier, our oligos have only two motifs concatenated by a spacer. Thus, we designed our eight bridge oligos to double in role as adapters that will include an adenosine overhang after annealing to address(*A*_0_) and payload(*P*_0_) oligos (Fig 3.b). The address motifs are 5’ phosphorylated which results in all oligos in our pool having their 5’ end analogous to end-prepared dsDNA. Thus, these oligos can readily ligate with the AMX sequencing adapters from ONT’s ligation sequencing kit (LSK-109). The AMX adapters were attached to the oligos in a 10 minute reaction. Sequencing was performed on a R9.4.1 flow cell for 4 hours. Basecalling was performed with both Guppy and Bonito basecallers. The sequencing run generated 27,198 reads with an N50 of 192bp.

Despite having several reads, we found that the reads were low quality. From the read length distribution in Fig 4.a and Fig 4.b, we see that the median read length with Guppy and Bonito is 166nt and 110nt. Thus, more than half reads are 48% longer than original oligos as several reads were observed to contain multiple oligos in a single read. On further analysis, we identified wrong event detection by MinKNOW to be the root cause of the problem. When sequencing oligonucleotides on an ONT R9.4 flowcell, the movement of bases through the pore leads to a continual change in current, known as the “squiggle”, that is recorded by MinKNOW. MinKNOW processes the squiggle into reads in real-time, and each read is supposed to correspond to a single strand of DNA. However, as our oligos were below 200 bases, we observed that sequencing our oligos generated low quality reads due to incorrect segmentation by MinKNOW which would earmark empty signals as valid reads, and created reads with merged squiggles for more than one strand of DNA.

**Fig. 4.**
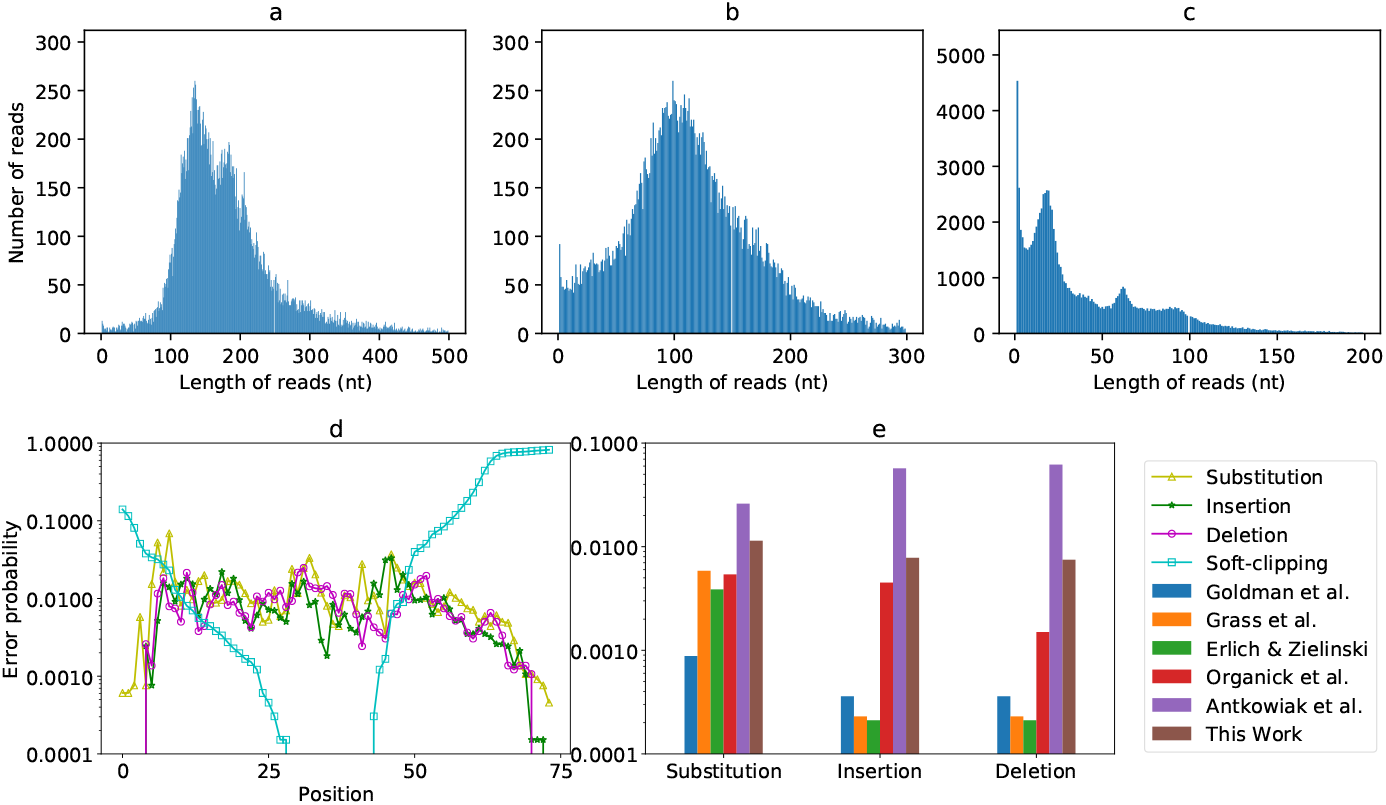
Analysis about the sequenced reads. (**a**) Read length distribution with Guppy basecaller. (**b**) Read length distribution with Bonito basecaller. (**c**) Read length distribution with Bonito basecaller post-processed with SaberSplit. (**d**) The substitution, insertion, deletion and soft-clipping rate per position of Guppy reads. (**e**) Comparison of errors rate with previous work.

Due to the presence of multiple oligos per read, we cannot directly align the reads to the reference oligos. So we did reverse alignment to study error characteristics and coverage distributions. We regard each read as a “reference” and build an index per read. Then, we treat each oligo like a “read”, and align it to each reference. Thus, for each read, we get an alignment file that contains one record per oligo. To identify and retain only good alignments, we filter the alignments using the following criteria: (i) MAPQ *>* 10 (90% alignment confidence), ii) all alignments in a read should correspond to one orientation (no mixed forward and reverse alignments), and iii) there should not be any overlap when multiple oligos are mapped in a single read; only the alignment with the highest alignment score is kept if several alignments overlap each other. With this approach, we get the set of oligos that we can identify assuming we have full knowledge of the original oligos.

Using Minimap2 [18] for reverse alignment, we computed the substitution, insertion, deletion and soft-clipping rate per position (Fig 4.d). As can be seen, the rate of soft clipping is very high at the extremities (especially 3’ end)due to the very high error rate caused by BOA and DOS. In the middle portion of the read, the rates of error types vary, with no one error type being dominant over others. These results are in sharp contrast to error statistics published in prior work on DNA storage [5, 7, 10, 19, 20], where substitution errors have been shown to be more likely than indel errors, and overall error rates are at least 10× lower (Fig 4.e and Supplementary Table 1). The only exception is work on photolithographic synthesis [21], where the error rates reported were also high.

**Table 1.** Processing time (in second) for real dataset with 12 CPUs

|  | 6× | 13× | 20× | 27× | 34× |
| --- | --- | --- | --- | --- | --- |
| Motif search exec. time | 0.15 | 0.24 | 0.38 | 0.49 | 0.64 |
| Reverse alignment exec. time | 29 | 60 | 91 | 122 | 152 |

### 2.4 Correcting Event Misdetection with SaberSplit

In the real DNA storage scenario, the original reference oligos must be inferred from erroneous reads automatically. Current read clustering and consensus callers used for this purpose assume that a read covers only a single oligo. To be able to use them, we developed SaberSplit (Supplementary Note 1), a tool that reduces the errors caused by incorrect segmentation by splicing squiggles to separate out reads belonging to different oligos. With SaberSplit, the original reads are chopped to 102,221 shorter reads of median length 25nt as shown in Fig 4.c. Then, we tried to use state-of-the-art clustering programs and position-wise consensus callers [22, 23] to infer the original oligos from both raw Bonito/Guppy reads, and SaberSplit processed reads. However, due to the high error rate, no oligos could be inferred in all cases.

To study SaberSplit further, we aligned the chopped reads to reference oligos with Minimap2. We compared the alignment statistics for raw Guppy, Bonito and Sabersplit processed reads (Supplementary Table 2). Guppy reads produced the highest number of alignments, with 102% more reads being aligned than Bonito. This could be explained by the fact Bonito is optimized to work better with longer reads, making it less suitable for short ones. Surprisingly, SaberSplit performed the worst with 9.5% fewer reads than even Bonito. This showed us that splitting reads amplifies the error rate and makes the case for a consensus caller that can directly work with raw reads covering multiple oligos.

### 2.5 Inference and Consensus with Motif Search

To reconstruct the original data from noisy reads, we developed a new reconstruction algorithm called Motif-Search that meets two requirements: (i) guarantee successful recovery despite high error rate, and (ii) directly work with raw, basecalled, Nanopore reads that might contain multiple oligos per read. Motif-Search differs from prior consensus callers that it is structure aware—while other callers view an oligo as a random collection of nucleotides, Motif-Search exploits the fact that our oligos are a collection of payload motifs separated by spacer motifs, with all motifs being drawn from a predefined, finite library. A detailed description of the Motif-Search algorithm is presented in Section 4.3. Here, we present our analysis results that demonstrate the ability of Motif-Search to accurately infer original oligos.

Fig 5 shows the true positive (TP) count (number of inferred oligos that are in the original set) of Motif-Search and Minimap2-based reverse alignment method at various coverage levels (lower sequencing coverage simulated via subsampling reads). It is important to note that Minimap2 needs the original oligos which would not be available in the real DNA storage use case. Thus, Minimap2 results are used as a baseline for comparison rather than a real decoding solution. First, Motif-Search is able to fully recover all oligos at 20× coverage. Reverse alignment misses one oligo even with 34× coverage. Second, Motif-Search reconstructs more oligos than reverse alignment at all coverage levels. The under-performance of reverse alignment relative to Motif-Search is because all the reads covering the missing oligo had a very poor alignment and were filtered out.

**Fig. 5.**
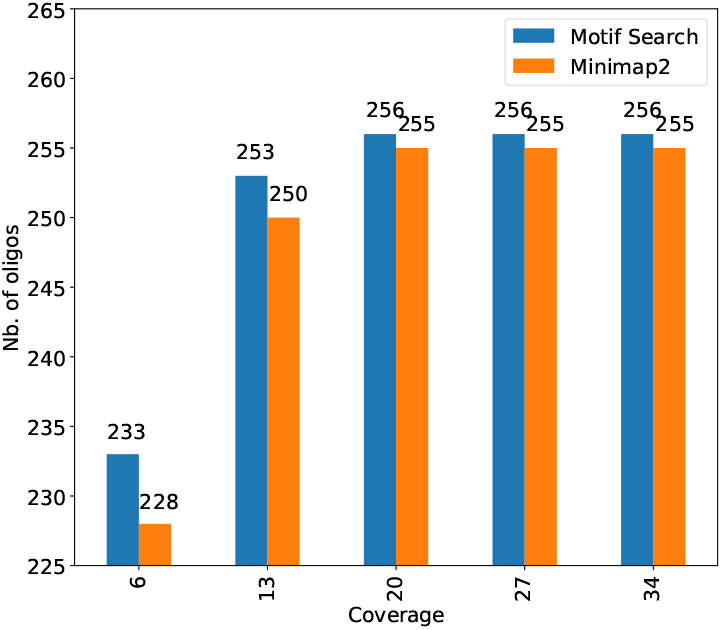
Number of oligos correctly reconstructed. Motif-Search fully recovers all oligos at 20× or higher coverage. Minimap2 misses one oligo even with 34× coverage.

Table 1 shows the execution time of Motif-Search and reverse alignment. Both support multi-threaded operation. On a 12-core Intel(R) Core(TM) i9-10920X CPU clocked at 3.50GHz, 128GB RAM with a 1TB SATA SSD, Motif-Search is 190–250× faster than Minimap2 due to the fact that Minimap2 needs to build an index for each read and align each oligo to each read while Motif-Search is custom-designed for the motif-based oligo reconstruction use case.

In order to investigate false positive (FP) behavior of Motif-Search and reverse alignment, we increase the motif library size. For a given set of address and payload motifs, we create oligos containing all possible combinations of motifs. For instance, if the motif set size is 64(*address*) × 256(payloa*d*), we generate 16,384 possible oligos. We then use Minimap2 to align each oligo to each read. We use the same reads as before which were sequenced from 256 original oligos. As the motif set is expanded, Motif-Search can now report an inferred oligo which is not in the original set but from the expanded set, which would be labelled a FP.

Fig 6 shows the TP and FP counts for various expanded motif sets. First, note that Motif-Search is able to reconstruct all original oligos when sequence coverage reaches 27× for all motif set sizes. When the sequence coverage is low, Motif-Search is able to reconstruct more true positive oligos than reverse alignment even though it is unaware of the reference oligos. Second, as the motif set size increases, the number of FP for both approaches rise. Since the sequences are error-prone, both approaches make errors identifying the correct references from reads. However, the FP rate of Motif-Search is still lower than reverse alignment. While missing TP is an issue as it can lead to data loss, extra FP is not a problem as it can easily be discarded by using auxiliary metadata and/or error-control coding.

**Fig. 6.**
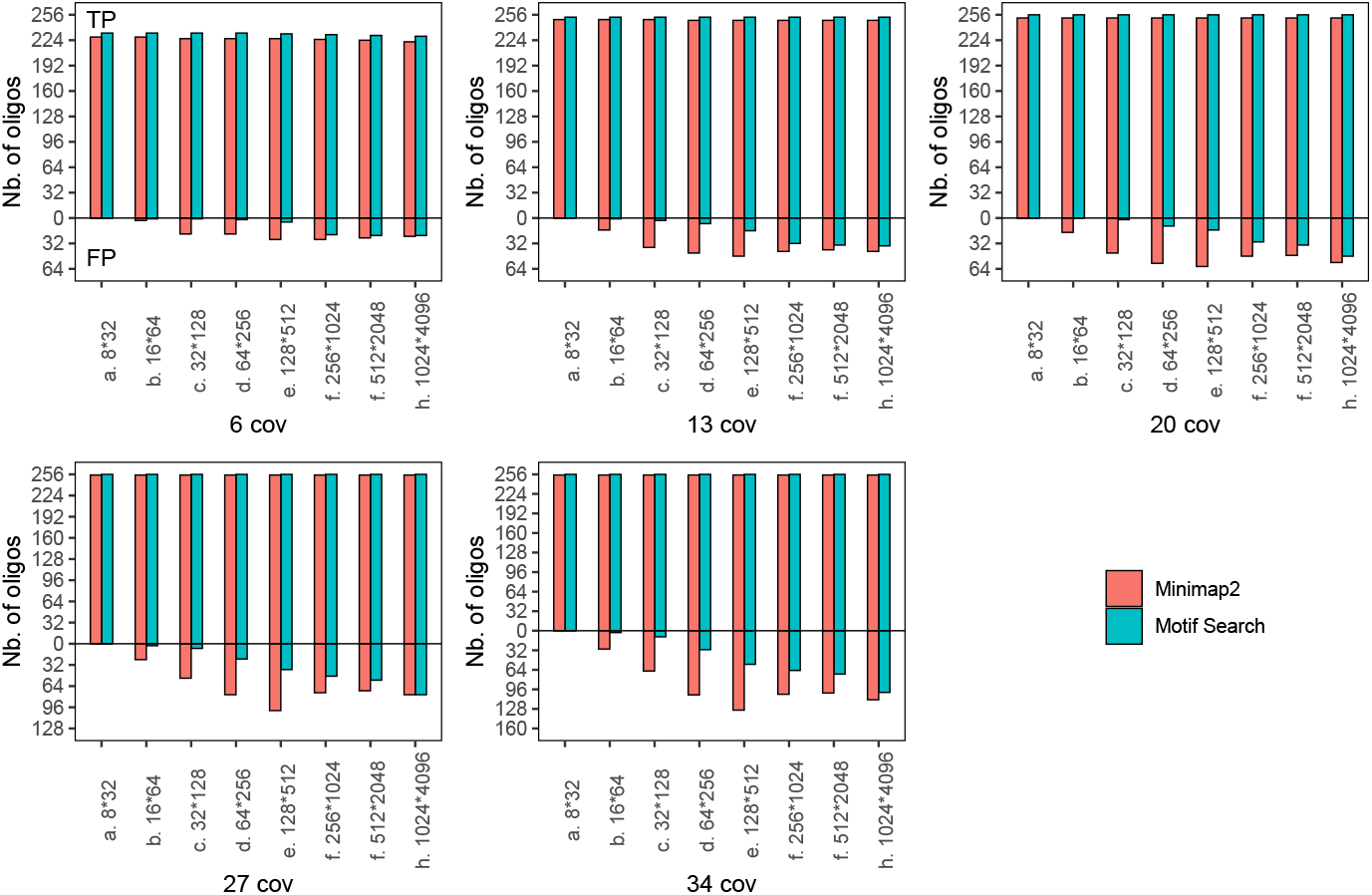
The number of true positive and false positive oligos reconstructed by Motif-Search and Minimap2 for different sequence coverages with expanded motif sets. i) Motif-Search reconstructs more true positive oligos than reverse alignment even without the knowledge of reference oligos. ii) False positive rises for both approaches when the motif set size increases.

These results clearly demonstrate that (i) our motif-based, BOA method can successfully encode information in DNA, and (ii) with sufficient coverage, Motif-Search is capable of reconstructing all original oligos, and thereby ensuring successful decoding, despite errors introduced by enzymatic BOA and DOS.

### 2.6 Read–Write Cost Comparison

The cost of storing data on DNA comes from two aspects, namely, the cost of sequencing for reading data and the cost of synthesis for writing data. Composite motifs has the potential to reduce the synthesis cost, thanks to the increase in logical density. For example, each synthesis cycle encodes 84 bits (*log*_2_*C*(96, 32)) in our composite motif experiment. A native motif-by-motif approach, in contrast, can only encode 6 bits per cycle with the same 96 motifs, and the traditional phosphoramidite approach can encode 2–3.37 bits per cycle depending on whether standard or degenerate bases are used for encoding. This 14–42× increase in logical density will lead to a proportionate reduction in synthesis cost over conventional synthesis approaches, as fewer synthesis cycles and fewer oligos are required to encode the same digital data. Since current motif-based synthesis techniques already use a high degree of sequence multiplicity, composite motifs can be easily integrated by generating a variant motif mixture pool without much added costs. The physical density of our approach is 3.36bits/nt, which is also higher than the physical density of conventional base-by-base DNA storage solutions (2 bits/nt) and comparable to degenerate base approaches (3.37 bits/nt[12, 13]).

While our solution improves logical density and synthesis costs, it does so at the expense of higher read costs. Fig 7 presents a comparison of the cost to read 1MB of data stored in DNA of our approach and other related work[5, 7, 10, 19, 21, 24] based on the cost of DNA sequencing (0.006$ per megabase) reported by National Human Genome Research Institute (NHGRI) in August 2021 [25]. The detailed calculation is included in Supplementary Note 2. Clearly, our work increases read cost compared to prior work except Antowiak et al. This is expected, as these prior approaches to DNA storage are able to fully recover the data at much lower sequencing coverages of 5× and 10× due to (i) the use of low-error rate array synthesis and high-throughput sequencing with extensive library preparation, and (ii) the use of error-control coding. Our current work, in contrast, focuses on (i) tolerance to errors introduced by enzymatic ligation and direct sequencing without any library preparation, and (ii) complete recovery without additional error correction. As synthesis is approximately 80,000 times more expensive than sequencing [7] and the read cost continues to drop due to rapid advances in sequencing, we believe that it is more important to focus on reducing the write cost, which is a bottleneck today in DNA data storage.

**Fig. 7.**
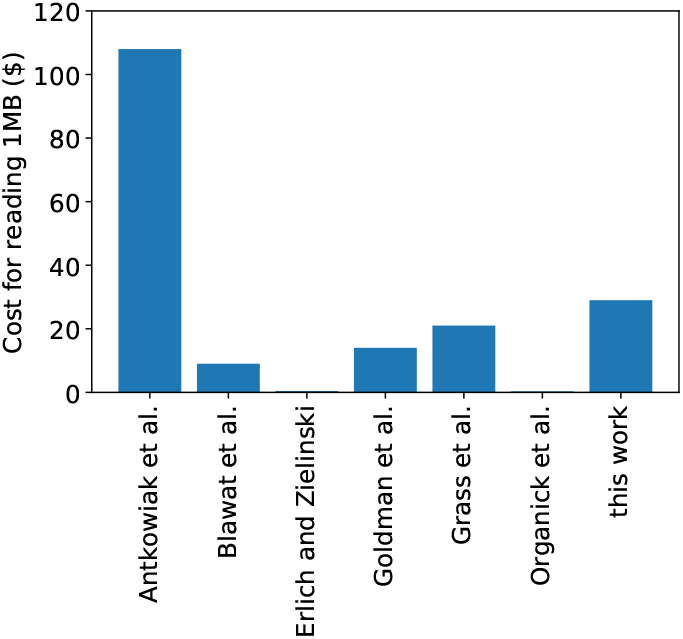
The cost of DNA sequencing to read 1 megabyte data. Our work increases read cost compared to prior work except Antowiak et al [21].

## 3 Discussion

In this work, we demonstrated the feasibility of using composite motifs to scale the logical density of DNA storage by an order of magnitude. We developed synthesis (BOA) and sequencing (DOS) methods customized for writing and reading oligos that regard composite motifs as building blocks, and showed that the error characteristics of these methods are different compared to state-of-the-art techniques. We developed a new motif-based consensus calling and oligo inference method (Motif-Search) that is able to recover all data at coverage as low as 20×. Our future work aims to scale up the methods presented in this paper on several fronts. First, to simplify the task of motif design, we built on an existing library of 25nt primers leading to a physical density of 3.36bit-s/nt. Future work will improve this further by optimizing the motif library. Second, we are working on reducing sequencing costs by adding error-control coding optimized to our DNA storage channel to enable data recovery at a lower sequencing coverage. Third, the short size of motif library, the library-preparation-free sequencing provided by DOS, and the error-tolerant nature of Motif-Search all simplify end-to-end automation. Thus, we are developing a fully automated DNA storage solution that can scale both oligo length and number of oligos beyond what was presented in this work.

## 4 Methods

### 4.1 DNA Assembly

Oligo with a format of *A*_0_–*P*_0_ (Fig 3.b) was realised with (i) a set of 8 ssDNA oligo sequences of 24-bases in length, representing *A*_0_; and (ii) a set of 32 ssDNA oligo sequences of 50-bases in length, representing the common spacer motif and each *P*_0_ motif. The sequences of motifs in these oligos were selected from 25mer DNA barcodes. A set of 8 ssDNA oligo sequences of 50-bases in length were designed to function as (i) a bridge between *A*_0_ and *P*_0_ for ligation; and (ii) an adenosine overhang on the 3’ end to facilitate AMX sequencing adaptor ligation.

#### 4.1.1 Phosphorylation

A pool of 32 oligos, representing the common spacer motif and each *P*_0_ motif, were 5’ phosphorylated using T4 PNK at a pool concentration of 300 pmol and reaction scale of 50 uL, as per the vendor guidelines at 37°C for 40 minutes. A denaturation step was performed to stop the phosphorylation at 65°C for 20 minutes.

#### 4.1.2 Assembly

The 8 *A*_0_ oligos and 8 Bridge oligos are pooled at equimolar concentrations and diluted to 25 uM final pool concentration. DNA assembly reaction was carried out by taking (1) 2 ul of the *P*_0_ phosphorylation mix, (2) 0.5 ul of the *A*_0_ and bridge pool (12 pmol) and ligated using (3) Blunt/TA master mix as per vendor guidelines. The *P*_0_ phosphorylation mix is composed of (1) 5 ul T4 PNK Rx Buffer, (2) 5 ul ATP (10 mM), (3) 1 ul T4 PNK, (4) 36 ul NFW, (5) 3 ul *P*_0_ (300 pmol). The above reaction is incubated at 95°C for 3 minutes and gradually cooled to room temperature.

### 4.2 Nanopore Sequencing

Sequencing sample preparation was carried out using LSK-109 kit. AMX sequencing adaptors were ligated by mixing 2.5 ul of the assembly mix with 5 ul AMX and 5 ul Blunt/TA mastermix from NEB and incubated for 10 minutes. The sample was then loaded into a R9.4.1 MinIon flowcell and sequenced for 90 minutes. Basecalling was performed on the Guppy (v4.0.15).

### 4.3 Motif-Search Algorithm

Motif-Search works in two stages, *inference* and *consensus calling*. In the inference stage, it maps each read to an inferred oligo. During consensus calling, it uses all inferred oligos to produce a consensus set of inferred reference oligos.

#### 4.3.1 Inference

The first task performed by Motif-Search is to extract one or more oligos from each read. Recall that an oligo is a set of motifs concatenated by spacers. Motif-Search infers oligos by first locating the spacer positions and then mapping the portions of the read between two spacers to the reference motifs to determine the payload and address motifs. Inference works in three steps: i) segmentation to locate spacer positions, ii) mapping to identify reference motifs between spacers, iii) overlap check to extract only oligos that do not overlap with each other.

##### Segmentation

Segmentation determines the spacer positions. Since all spacers are identical, their candidate positions can be located by k-mer seeding. We convert A, T, C and G into a two-bit equivalent representation and build the index of the spacer by extracting all k-mers of length four (found to be optimal experimentally). To process each read, we extract all 4-mers in the read, lookup the index, and collect positions with an index hit. The positions are adjusted by the offset of the k-mer to get normalized positions.

To eliminate candidate positions with low confidence, we filter out the positions having less than *spacer length/k* k-mer votes. As reads are error prone, indels can cause candidate positions that should be identical to differ slightly by a few nucleotides. This could result in candidates receiving fewer votes and failing the filter. Hence, we merge neighboring positions and represent them by a centroid with a combined count. At the end of this stage, we have all candidate positions for all spacers in a read.

In our experiment, each oligo has only one spacer. But in the general case, each oligo can contain multiple spacers. From the structure of the oligo, we know that each oligo with *M* motifs has *M* − 1 spacers, with each spacer being spaced apart by a distance *d* equal to the sum of the motif length and spacer length. In order to accommodate synthesis and sequencing errors, these interspacer gaps can be slightly more or less than the motif length depending on indel errors. Thus, we identify all possible chains of *M* − 1 positions which are within an expected distance threshold from each other.

As mentioned earlier, the candidate positions in these chains are approximate, as indel errors can result in observed starting position differing from actual starting position by a few nucleotides. We rectify and refine these positions to tolerate indel errors by using randomized embedding—a technique which has been demonstrated to be a scalable approach for mapping reads to references in genomic sequence alignment[26]. More specifically, for each candidate position, we extract a spacer-length portion of the read at that position and at several positions around that position. We embed each extracted read fragment using a randomized algorithm and compare with the embedded version of the original spacer motif using hamming distance. We select the shifted position with least embed distance as the final candidate position. As the number of candidate positions can be large, the use of embedding helps us to avoid expensive edit distance computations between the read and spacer motif, and use hamming distance between their embedded versions to rectify candidate positions.

##### Mapping

Given a chain of refined candidate positions, we can extract the potion of each read between two neighboring spacers. These portions correspond to address and payload motifs. The next step is to identify the original motif for each observed motif in the read. This can be translated to a sequence mapping problem by considering the original motif library as the reference and the observed motif in the read as the query. Therefore, we use the ksw-lib([27]) to select the optimal original motif with the highest mapping score for each observed motif. After this step, we have multiple chains of mapped motifs.

##### Overlap check

As we consider all possible chains, some chains might overlap each other. However, while each read can cover multiple oligos due to DOS, each nucleotide in a read should map to only one motif/oligo. Thus, the final step in the inference stage is to identify the optimal set of chains that do not overlap with each other. To do this, we traverse the chains to identify overlapping sets. For each overlapping set, we pick a chain with the highest mapping score such that no chain appears in two sets.

#### 4.3.2 Consensus Calling

Each original encoded oligo can be synthesized with duplication. Library preparation steps, like PCR, also amplify the pool of oligos by creating multiple copies of each oligo to ensure successful sequencing. Thus, an original oligo can be covered by multiple reads. For each read, the inference stage identifies the optimal set of non-overlapping chains. As the final step, we apply consensus calling to group similar motif chains inferred from the *inference* stage, and obtain consensus to achieve higher confidence. We do this by first clustering the inferred oligos using their address motifs. Then, we select the most frequent motifs at each position as the final consensus motif as shown in Fig 8.

**Fig. 8.**
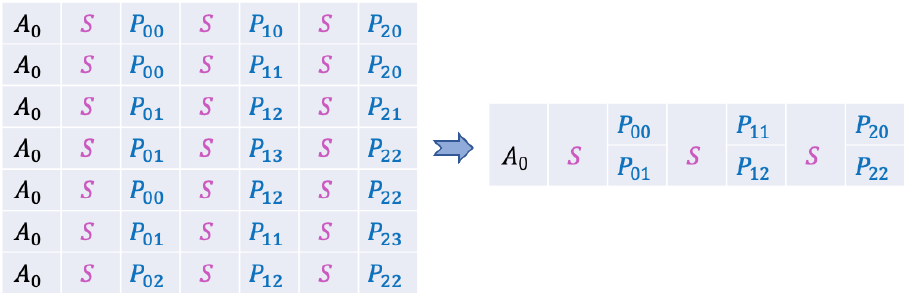
Example showing consensus calling with seven inferred oligos with the same address motif *A*_0_. The payload motifs are decoded as *P*_00_, *P*_01_ at the first position, *P*_11_, *P*_12_ at the second position and *P*_20_, *P*_22_ at the third position which are the *topN* (N is the number of oligos in each sequence) frequent motifs in each column position.

## Declarations

### Data availability

The oligo sequences are available via https://drive.google.com/file/d/1kf1XmU7cP3GNb1obdZravhFEPaZdu2A/view?usp=sharelink. The reads with Guppy basecalled, Bonito basecalled and Bonito basecalled post-processed by SaberSplit are available via https://1drv.ms/u/s!AuZMWsmSlzpWqvYSb43qImGaJ4haA?e=FhoZuz, https://1drv.ms/u/s!AuZMWsmSlzpWqvYVM3S6gaZR33FRwA?e=Te6naa and https://1drv.ms/u/s!AuZMWsmSlzpWqvYO1I7upDWuujWaiQ?e=53Fgv3.

### Code availability

The Motif-Search algorithm implementation is available via https://gitlab.eurecom.fr/yan1/motif-search under MIT license. SaberSplit is available via https://github.com/helixworks-technologies/sabersplit.

### Author contributions

N.P., S.C., and C.C. developed and performed the wet lab experiments. Y.Y. and R.A. performed algorithm design, software implementation and simulations. R.A., Y.Y. and N.P. analyzed the experiments and wrote the paper. All authors read and approve the final manuscript.

### Additional information

The supplementary information is attached.

