## Supplementary for "Scaling Logical Density of DNA storage with Enzymatically-Ligated Composite Motifs"

### SUPPLEMENTARY INFORMATION

**Supplementary Table 1.** The substitution (SUB), insertion (INS) and deletion (DEL) rate of SOTA work. Data for Goldman et al. [1], Grass et al. [2], Erlich Zielinski [3], Organick et al. [4] and Antkowiak et al. [5] was taken from Antkowiak et al. [5].

| | Goldman et al. | Grass et al. | Erlich et Zielinski | Organick et al. | Antkowiak et al. | This Work |
|---|---|---|---|---|---|---|
| SUB | 0.00088 | 0.005850 | 0.003870 | 0.005400 | 0.026000 | 0.011411 |
| INS | 0.00036 | 0.000230 | 0.000211 | 0.004500 | 0.057000 | 0.007817 |
| DEL | 0.00036 | 0.000230 | 0.000211 | 0.001500 | 0.062000 | 0.007485 |

**Supplementary Table 2.** Statistics of the reads.

| | Nb. reads | Nb. aligned reads | Median read length |
|---|---|---|---|
| Guppy | 27198 | 9960 | 166 |
| Bonito | 27198 | 4901 | 110 |
| SaberSplit | 102221 | 4434 | 25 |

**Supplementary Note 1.** Sequencing data and SaberSplit.

The generated sequences via the BOA method were sequenced via DOS for 4 hours to generate 27,198 reads with an N50 of 192bp. The reads were basecalled using guppy basecaller (v4.0.14) in high accuracy mode. We suspected that some of the reads would not be split properly by the MinION instrument as observed in prior research. To split the concatenated reads and to improve basecalling, we developed a nodejs script called SaberSplit to correctly identify the adapter regions of the reads, which has an unique pA (picoamperage) signature that can be identified from the events data of the fast5 file.

SaberSplit is a node.js based tool to process the .TSV files generated from the SquigglePull program of the SquiggleKit(`https://github.com/Psy-Fer/SquiggleKit`). SquigglePull TSV files contain the read-IDs and their event level data in the following format. SaberSplit extracts the event data from the TSV files and stores them in an array. It calculates the median and MAD (Median Absolute Deviation) of the event data. It calculates (Datai-Median)/MAD for each of the data points and if the (Datai-Median)/MAD > 5. It takes that data point for further processing.

SaberSplit extracts the events on the right-hand and left-hand side of the triggered event if they have (Datai-Median)/MAD > 3. If the total number of extracted events that are on the left and right along with the triggered event is less than 12 events. The event is classified as a spike and the read is split. SaberSplit processed 27,198 reads, generating 237,327 new split reads. Many of the split reads are short and have not passed the threshold for basecalling with bonito. A total of 102,222 were successfully basecalled from the SaberSplit reads. Guppy generated an extremely low number of successful basecalled reads from the SaberSplit reads.

**Supplementary Note 2.** Sequencing cost projection.

- The sequencing cost to read 1 Megabyte is simulated as the Equation 1.

$$reading\_cost = oligo\_length * nb\_reads * sequencing\_cost\_per\_nt / stored\_data\_size \quad (1)$$

- The oligo_length includes the length of primers.
- The sequencing_cost_per_nt takes the value 0.006$ per megabase reported by National Human Genome Research Institute (NHGRI) in August 2021.
- Data for Goldmann et al. [1], Grass et al. [2], Erlich et al. [3] and Organick et al. [4] was taken from Organick et al. [4].

Table 1: Sequencing cost projection

| | oligo length | nb of reads | data size | sequence cost ($/MB) |
|---|---|---|---|---|
| Antkowiak et al. | 60 | 30000000 | 99103bit | 108 |
| this work | 74 | 640 | 80 bit | 29.8 |
| Grass et al. | 159 | 1858027 | 679000 bit | 21.9 |
| Goldman et al. | 183 | 7960000 | 5200000 bit | 14.1 |
| Blawat et al. | 230 | 144475005 | 22MB | 9.06 |
| Erlich and Zielinski | 200 | 750000 | 2.11 MB | 0.43 |
| Organick et al. | 150 | 67241860 | 200MB | 0.3 |
